# Using Fiji as a simplified tool for assessing alveolar chord length and septal thickness in neonatal pulmonary disease research

**DOI:** 10.1101/2025.05.29.656796

**Authors:** Pragnya Das, Emanuele Martini, Osama Aloudat, Emily Tran, Divya T Rajendran, Beamon Agarwal, Robert J Homer, Vineet Bhandari

## Abstract

Bronchopulmonary Dysplasia (BPD) is a neonatal condition primarily affecting babies born prematurely, who have been exposed to invasive ventilation and/or oxygen supplementation. These therapies predispose the immature developing lungs to inflammation and injury to the alveolar epithelium resulting in loss of the alveolar epithelium and thickening of alveolar septa, the 2 important hallmarks of the pathology of BPD. The expansion and enlargement of the alveolar sacs gives the characteristic feature of alveolar simplification in BPD. To measure this alveolar abnormality, we have modified and developed a custom-made Chord Length Fiji plugin which can easily measure the Chord Length and Septal Thickness on Hematoxylin-Eosin-stained histological lung sections, thus generating an instant morphometry readout for quick interpretation. We validated this plugin in multiple animal experimental models and human neonatal lungs with BPD. We were able to differentiate the morphology under conditions of ventilation or hyperoxia exposure, when compared with untreated controls. We thus conclude that our Chord Length and Septal Thickness plugin can be used in a simplified manner following a less meticulous automated operation. To facilitate global use, we have created a DOI so that any user can access it without any restrictions to complete the morphometric measurements. This plugin will be useful for smaller research laboratories with limited budgets and require no access to complex software.

## Introduction

Morphometry allows objective quantification of the shape of any biological structure. As such, it is an indispensable component of the statistical analysis of size and shape variation in biological structures and can be applied using computer vision techniques. In lung research, it is essential that we understand the quantitative anatomy of the lung and its association with pathology and physiology because different abnormalities in the lung secondary to varied disease conditions can result in a typical alveolar structure, characteristic of that specific disease. This change in shape of the lung can be morphometrically assessed using the principle of stereology wherein unbiased sampling and different measurement tools are needed to estimate total lung volume, volume of lung parenchyma, alveolar surface area and number of alveolar epithelial cells per lung [1].

The history dates back to the pioneer pulmonology researcher, Ewald R Weibel who was the first to define “morphology” as the link between genetics and function in the context of lung biology [2] and introduce morphometry in lung research [3] by employing light microscopy to study finer details in lung structure. With the help of his colleague, Domingo M. Gomez, together they developed mathematical algorithms to estimate alveolar abnormalities in different pulmonary malfunctions. Since then, morphometry has been used as a powerful tool to measure elements of pulmonary architecture in various pulmonary diseases and conditions such as chronic obstructive pulmonary disease (COPD), emphysema, fibrosis, respiratory distress syndrome (RDS) and bronchopulmonary dysplasia (BPD). More importantly, we would also like to make the readers aware of the immense contribution of Wayne Rasband [4] who wrote this Program and developed it under the name “NIH Image” during his tenure as a software developer at the Research Services Branch (RSB) of the National Institute of Mental Health (NIMH), part of the National Institutes of Health (NIH, USA). It was later superseded by ImageJ followed by ImageJ2 and is now operational as “Fiji”. John C Russ, a faculty in the Materials Science and Engineering Department of North Carolina State University (NCSU), then recorded a set of macros and plugins in ImageJ for image processing [5, 6], which was later considered a gold standard for measuring various biological shapes and structures. Mr. Rasband retired from the NIH in 2010 but continues to support and monitor the advancements in the software as a “special service volunteer”. Dr. Russ has also retired from NCSU and continues to consult various academic institutes, defense and federal labs through Reindeer Graphics.

Traditionally, morphometrics has always relied on specialized software like .tps series [7] and “R” packages [8-11] which incorporates computer vision into the software and most computer vision libraries are implemented in dlib using Python or C++ [12]. Automated stereology improves accuracy with less effort as compared to manual morphometry. Recently software such as Deep-Masker [13] STEPanizer [14], newCAST^TM^ (from Visiopharm [15], STEREO INVESTIGATOR (from MBF BioScience) [16], STESYS [17, 18], STEREOLOGER (from SRC BioSciences) and STERapp (from STEREOLOGY TOOLKIT) have become more popular for automated image quantification as compared to some traditional image analysis tools like CellProfiler (http://www.cellprofiler.org/), iCluster (http://icluster.imb.uq.edu.au/), wnd-charm (http://code.google.com/p/wnd-charm/) and PhenoRipper (http://mondale.ucsf.edu/Web_Site/PhenoRipper/default.htm) [19]. With modern resource constraints, then free morphometry software like Fiji [20] (the updated version of ImageJ created by Johannes Schindelin and currently maintained by the Laboratory for Optical and Computational Instrumentation at the University of Wisconsin in Madison, USA) for global academic use, is an attractive alternative. The major barrier to widespread use is the generation of plugins from reliable secure sources. We wanted to make available a Fiji plugin for Chord Length assessment in a simple, reproducible, user-friendly approach. The aim was to generate uniform data, globally, especially for those working in neonatal lung biology that involves morphometric measurements. Hence, in the present work, we will describe the function of this macro and the steps to download, install and use the relevant macros to measure Chord Length (and Septal Thickness) in the experimental animal models, and compare it with normal controls.

### Lung harvesting, tissue sectioning/staining and image acquisition

Only 3 tools are needed to evaluate the Chord Length and Septal Thickness - well perfused lungs, good histological Hematoxylin-Eosin (H/E) stained sections without any interfering blood vessels or airways, and sharp, pristine photomicrographs taken at 100X total magnification. For the mouse lung, to get the best histological lung sections, during harvesting of tissues, the heart should be perfused with cold 1X PBS (phosphate buffered saline) or freshly prepared cold 4% PFA (paraformaldehyde), and lungs inflated at a pressure of 25cm H_2_O to clear the organ of any residual blood clots. A well-perfused lung should look clean and translucent (**Fig. S1a**). Since it is critical to have uniform images for unbiased comparisons, it is important to acquire the best images for morphometry quantification. To do so, we suggest the use of the left lung only for histological purpose for the simple reason that the left lung has only 1 lobe while the right lung has 4 lobes. Hence, during microtomy sectioning of the right lung, empty spaces between each individual lobe may interject the entire lung which will confuse the software and interfere in the final read-out. Therefore, it is ideal to use the left lung because of its simple single-lobed structure. It is also advisable to section it from the dorsal side as shown in **Fig. S1a** (to avoid any intervening pulmonary blood vessels and capillaries). It is also important to collect the superficial sections rather than going deep towards the central region of the lung where there are abundant branches of the central pulmonary artery and the major central airway. Since the idea is to acquire images only with alveolar sacs *sans* blood vessels or respiratory bronchioles, sectioning the lungs from the ventral side is not a good idea. Deeper sections result in the inclusion of these large airways and blood vessels (**Fig. S1b**) which will add false values to the Chord Length readout. As Fiji can count any empty white space as an alveolar area and record erroneous data, the problem can be overcome by the above sectioning methodology without the hassle of masking empty spaces in lung sections either manually or digitally using different computer-generated programs. Similarly, staining is another important factor that must be considered prior to image acquisition. Poor staining as seen in **Fig. S1c-i** will confuse the software and the plugin will not be able to adequately overlay the image (**Fig. S1c-ii)** or convert it appropriately to binary mode (**Fig. S1c-iii)** for the final readout **(Fig. S1c-iv).** H&E staining on the tissue sections should be crisp and clean as seen in **Fig. S1b.**

### Morphometry in the context of BPD (experimental as well as clinical)

BPD is a neonatal respiratory condition that predominantly affects infants born prematurely ≤28 weeks of gestation. Premature infants susceptible to develop BPD are therefore born in the late saccular to the early canalicular phase of lung development [21, 22], a stage when the terminal airways start to differentiate and vascularize, and surfactant production is initiated from the Type II alveolar epithelial cells. The etiology of BPD is multi-factorial [23]; oxygen-related toxicity is one of the most important contributing factors to impaired alveolarization characterized by the simplification of the alveolar epithelium resulting in enlarged alveolar spaces. Hence, it is important to measure this surface area to compare the degree of abnormality with normal developing lungs of babies born full-term. Alveoli formation is a continuous process, which in humans starts between 32nd-36th week of gestation and continues postnatally up to 8 years of childhood [24, 25], whereas in mice it is initiated at P4 (postnatal day 4), peaks at P7 and ends around P36 [26]. In both cases, there is progressive transformation of the saccules to alveoli throughout the continuous phase of lung development wherein lung saccules divide constantly by outgrowing secondary septa, resulting in an increased number of smaller subunits called alveoli. This process of alveolar division is greatly impacted by different disease conditions. Consequently, the total alveolar surface area is considerably altered. Hence, alveolarization is dependent on pulmonary microenvironment that includes changes in at least three morphologic characters: alveolar size that impacts total alveolar volume, total alveolar surface area and total number of alveoli. The unique abnormality in BPD is that a developing lung is seriously impacted with incomplete morphogenesis due to disruption of definitive alveoli by secondary septation of primitive saccules. Upon exposure to hyperoxia, the alveolar sacs are injured resulting in a simplified alveolar epithelium and enlargement of the alveolar sacs, which leaves a damaging footprint throughout the life of BPD-affected infants. Due to the non-availability of a standardized method to measure this abnormality in larger animal models of BPD, researchers have used other parameters like radial alveolar count, distal airspace wall thickness, volume density of secondary alveolar septa, and branching morphogenesis [27-29]. We hope our customized Chord Length plugin will enable those researchers to measure this parameter in their BPD model as well.

### Comparison of Normoxia and Hyperoxia results in the context of BPD

The first line of identification in hyperoxia (Hyx) treated animal models is enlarged alveoli with thin, simplified alveolar epithelium (**Fig. 1a**) when compared with normal alveoli of the same age **(Fig. 1b).** This comparison is important because following Hyx exposure, we need to compare the structure of these Hyx-exposed alveoli (characteristic BPD pulmonary phenotype) with normal room air-control alveoli (who were *not* exposed to Hyx), with alveoli/lungs that may been treated with drugs to counteract the Hyx-induced lung/alveolar injury. This response is evaluated by measuring Chord Length as the primary visual morphological index of recovery to reflect significant structural changes that can be a focus of therapeutic approach, followed by molecular, biochemical, physiological, pathological and functional changes.

**Fig. 1:**
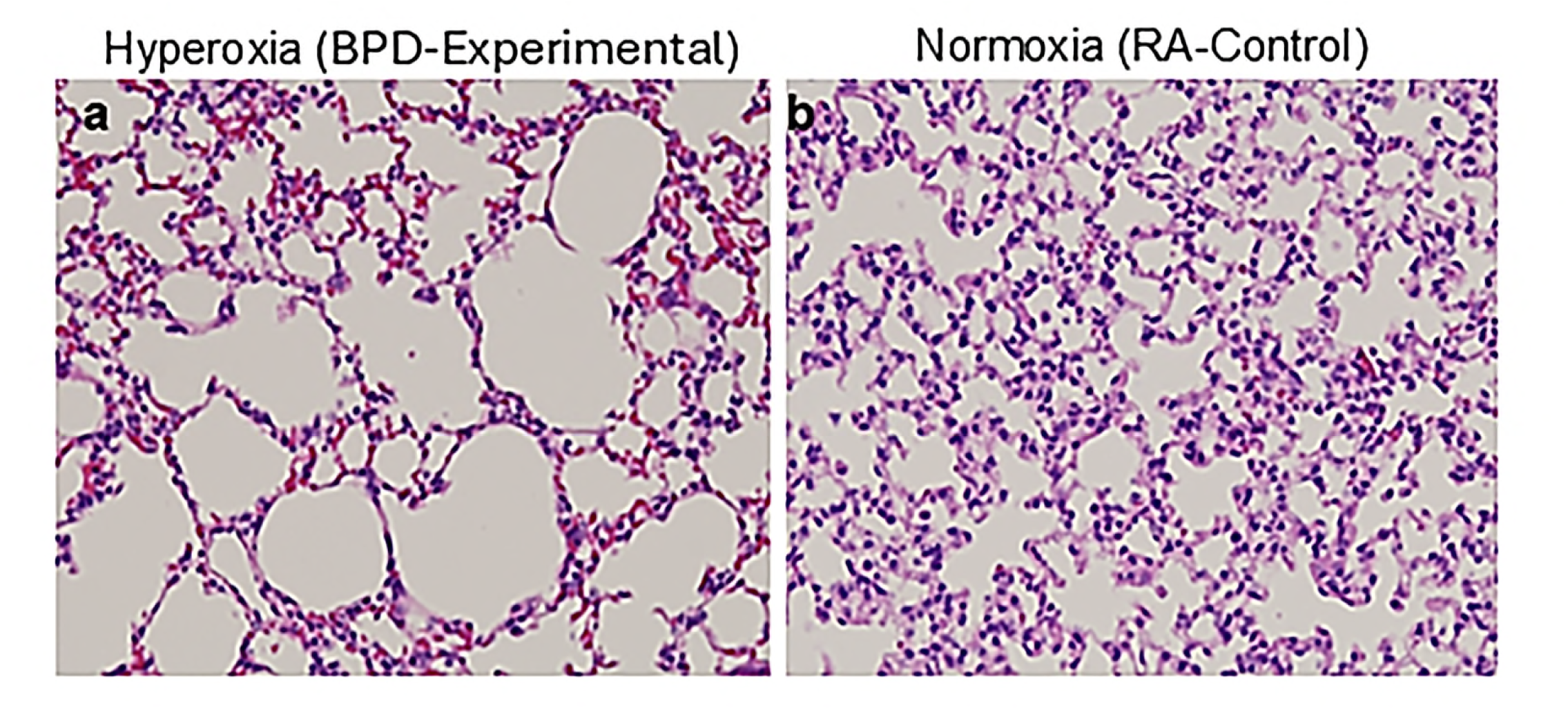
Histological section of P14 mouse lung showing **(a)** enlarged simplified alveoli in hyperoxia-exposed BPD group as compared to **(b)** control room air (RA) group. **P:** postnatal day; **BPD:** Bronchopulmonary dysplasia.

### Chord Length (L_m_) and Mean Linear intercept (MLI)

From the early sixties, ever since M.S. Dunnill introduced the concept of MLI to quantify emphysema in histological samples [24], MLI has been perceived as an index of airspace size, although formally it is a measurement of surface area-to-volume ratio. Later, another parameter i.e. Chord Length (originally denoted as L_m_; hereafter will be denoted as L_m_ for the readout and ‘Chord Length’ and/or ‘Septal Thickness’ when referring the plugin) was also introduced as a parallel surrogate for MLI. L_m_ is an indirect measure of alveolar size whereas MLI is a measure of the average distance (length) between alveolar walls [30]. Both serve as critical endpoints in animal models of BPD by providing an indirect estimation of the size and density of alveoli in lung tissue. Since MLI is a measure of morphometric change based on serial measurements of the lung using test lines, this parameter can reflect the average alveolar size and can be used to assess lung structure and function in various respiratory conditions but cannot be directly interpreted as a measure of alveolar size [31]. Simultaneously, the drawback of MLI is that since the parameter is based on hand-counting of intercepts from a limited number of representative histological sections, the grid lines must be drawn manually, the intercepts counted manually and then the final calculation also done manually, to estimate the value. In such a case, there is a lack of uniformity and possibility of a bias in the final read-out. While MLI comprises both the airspace and alveolar septum (**Fig. S2**), L_m_ only measures the ‘mean free path’ within the distal airspaces i.e. alveoli and alveolar ducts taken together (**Fig. S2**). Ironically, they are used interchangeably as synonyms in several publications; this is incorrect because a reported L_m_ value is always lower than the corresponding MLI value due to the fact that alveolar septal wall thickness contributes to the latter [32]. For e.g. the alveoli of a normal neonatal mouse lung at P14 would give a mean MLI score of 80µm [33], while the same alveoli will yield a mean L_m_ score of 40µm for the same mouse. Similarly for humans, while we found the mean L_m_ to be approximately 27µm for preterm controls, 40µm for term controls and 133µm for BPD patients, the MLI was found to range between 120-171µm for controls and 284µm for a BPD patient, as reported earlier [34]. However, there are certain other caveats, too. In some pathological conditions, there is increased thickness in the alveolar septum, with minimal or no change in the alveolar air space compared to the appropriate corresponding controls, while in others, the alveolar diameter increases with minimal or no change in the Septal Thickness. Similarly, if a researcher studies emphysema or fibrosis, it would be ideal to use MLI because both are associated with changes in lung elasticity and lung volume. On the other hand, in experimental BPD models, measuring L_m_ of the alveolar sacs is the recommended measurement, as L_m_ is a function of lung volume; in BPD the alveolar sacs are expanded with a simultaneous increase in lung volume [35]. Hence, L_m_ will not be able to differentiate the effects caused by tissue destruction *versus* tissue distension if used in conditions like emphysema or fibrosis. In such a scenario, the onus is on the end-user to decide what metric to use in the context of the study. We realized this shortcoming while going through the work of many researchers in the pulmonology field who had used MLI as an index to measure alveolar airspace; there were even some who quoted as using MLI as a measure of alveolar size. Although both terms are correct, it is important to know what morphological parameter should be used in the context of the disease studied rather than using a generalized format. It is therefore important to differentiate between “alveolar diameter” and ‘L_m_’ in the context of lung morphometry. While "alveolar diameter" refers to the full width of an individual alveolus (if one could directly measure it in 3D), ‘alveolar L_m_’ is a measurement taken across an alveolus on a 2D image (**Fig. S2**), essentially a straight line through the alveolus, which is usually smaller than the true diameter and is considered an indirect measure of alveolar size, particularly useful in studying lung diseases like BPD and emphysema where alveolar sizes are significantly affected.

Hence, our objective in writing this review is to introduce Fiji as a simplified software application for measuring L_m_ in experimental murine models of neonatal lung research, with a possibility of extending it towards large animal models and humans. Despite the abundant availability of online tutorials on different aspects of Fiji, there is not a single video or publication that highlights or cites L_m_ measurement in animal models using Fiji, from start to finish, though Artificial Intelligence (AI) may be able to overcome this limitation soon. Although Sucre Lab has attempted to design an AI based algorithm, the program measures MLI instead of L_m_ (https://github.com/SucreLab/AlveolEye). In sum, as L_m_ is currently not available in a single source publication, we have attempted to collate the information about L_m_ and compile it in an organized format to be presented as a more systematic scientific protocol for researchers in the pulmonary arena. We also introduce an additional parameter – Septal Thickness (S_T_) so that these dual parameters can be measured from a single image with our custom-made plugin. In the next section, we present a detailed step-by-step method to install Fiji with the specific plugin for L_m_ and S_T_, and measure both these parameters on the same histological sections, with a single click on the keyboard.

#### Septal Thickness (S_T_) as an indicator for BPD progression and other neonatal pulmonary diseases

As discussed earlier in this review, although oxygen supplementation and/or mechanical ventilation is currently a standard-of-care therapeutic approach for managing preterm babies, there is thickening of the inter-alveolar walls (septa) if these babies develop BPD. Consequently, gas exchange is significantly impacted causing respiratory difficulties [36]. Thus, increased S_T_ can also be counted as another hallmark feature of BPD pathology. The same holds true for animal models of experimental BPD, as well. In many knockout and transgenic neonatal mice models, deletion and/or overexpression of certain genes cause thickening of the septa [37-39] (as shown in **Fig. S2**), thus interfering in normal alveolar development and pulmonary function. Hence, it was important to include this parameter as a parallel index of morphometry along with L_m_.

### Organization of acquired images

Prior to launching Fiji, all the images to be analyzed should be systematically organized in respective folders names - **‘input’** (images to be analyzed) and **‘output’** (empty folder where all analyzed images with values will be stored) and should be saved in a master folder. In the present study, we have 5 groups organized in separate folders, each folder representing a separate species. Multiple lung images from each of these groups (with at least 3 animals from each group) should be acquired with the same magnification, using the same microscope and saved under the respective subfolders, as seen in **Fig. S3**. This organization is important because when calculating L_m_ and S_T_ in a batch for multiple images, the plugin will run through all the images in the folder thereby calculating the respective values in a single run.

### Installation of Fiji

Open your web browser and navigate to the official Fiji website https://fiji.sc/ where you will see download options for different operating systems. Click on the link that corresponds to your operating system (e.g., Windows, macOS, or Linux).

1. **Download the Installer:** Click on the appropriate download link for your operating system. This will typically start the download automatically with all the associated files (plugins, macros, etc.) in separate folders. Save the installer file to a location on your computer.
2. **Install Fiji:** Once the download is complete, locate the installer file on your computer and run it. Follow the installation prompts, including selecting installation options if prompted. The installation process may vary slightly depending on your operating system.
3. **Launch Fiji:** After the installation is complete, launch/run Fiji from your computer’s applications or programs menu.

A functional Fiji is now installed on your PC with all plugins attached to the program. However, a lot of times, additional plugins that are needed for your specific use may not be downloaded while downloading the original Fiji version. This is because the scripts for these additional plugins have not been written. In such a case, users may visit GitHub or Zenodo or any other publicly available software repository, designed for scientific sharing, and download additional plugins which are written in the same language as that of Fiji codes (Java, Python, Jython) for compatibility. To do so, close the Fiji application, and then download the required plugin from the necessary site. While installing, make sure to download this extra plugin in the Fiji installation directory into the “plugin” folder of the Fiji software saved in your PC. In the present work, we have modified an original Chord Length macro (Homer, unpublished script and publicly available; **SF1**) and a grid macro (originally scripted by John Russ and publicly available, **SF2**) and translated it in Python to be placed under the downloaded Fiji Folder, so that all plugins are housed in one folder in the same location. Any user wanting to modify the original script is free to do so with the intent of benefiting other users.

### Installation of the Chord Length ToolSuite

1. **Install the Chord Length ToolSuite** as shown in **Fig. 2a** by downloading first and then copying/moving it to the Fiji Plugin subfolder
2. **Launch Fiji and Set Background options** To do so, go to: **Edit** ➔ **Options** ➔ **Colors** ➔ and set <u>**Foreground:**</u> White <u>**Background:**</u> Black (Fig. 2b) **Process** ➔ **Binary** ➔ **Options** ➔ check □ **Black background (Fig. 2c)** The background option needs to be set only once. The ToolSuite is now installed with plugins for Single image and Multiple images analysis for Chord Length, Septal Thickness and all Other Alveolar Parameters. **Launch Chord Length & Septal Thickness – Single Image** (*abbreviated as Chord & Septal-Single Image*) and **Other Alveolar Parameters - Single Image** (*abbreviated as Alveolar Params-Single Image)* **Plugins** The input values for these 2 plugins are included within the description for ‘Multiple Images’ and are therefore not described separately (**Figs. 2d, f**) **Launch the Chord Length & Septal Thickness - Multiple Images** To do so, go to: **Plugins** ➔ **Chord Length ToolSuite** ➔ **Chord Length & Septal Thickness - Multiple Images** (*abbreviated as Chord & Septal-Multiple Image*) (**Fig. 2e**) Insert the input parameters in the tabs that pop with the following contents as described below: ***Input Directory***: Browse and select the folder that has all the images to be tested as described in **Fig. S3** ***Output Directory***: Browse and select an empty folder (duly named) which will save all the images after being analyzed as described in **Fig. S3** ***File extension***: .tif (if the images have been acquired in .tif format) ***File name contains***: keep this field empty (the program will analyze all the files in the folder. If the name of a single file is entered in the above fields, then the program will only analyze that individual file). ***Keep directory structure when saving:*** check this box □ to keep the same sub- directories organization of the input directory ***Pixel to µm calibration:*** We have entered this value as “0.7”. (*This value will vary for different microscopes and users should refer to Fiji/ImageJ user guide for appropriate instructions*). Entering this value will avoid calibration for all the images acquired and stored in the ‘input folder’; it is not necessary to acquire the images with the scale bar embedded within the image. This information is needed only once for a single image, for calibration purpose only. Failure to do so will require the photomicrographs to be captured with the scale bar embedded, and then the user must manually convert the distance of the scale bar (in pixel) to µm, every time. While acquiring the images, make sure all the images are acquired in the same magnification [e.g. 100X total in this study] using the same microscope, and the scale bar is not embedded in the image. Using an image with the embedded scale bar will interfere in the algorithm. ***Grid size for L_m_***: Usually 2 grids are made - one using only vertical lines and the other using only horizontal lines, the location of which are put in the appropriate part of an alveolus. For the present study, it is 32 pixels (this is the size of the grid used for the analysis of images in pixel with a total magnification of 100X. While calculating the grid size that enclosed and intersected a single alveolus, we found the number to be 32 (based on the macro scripts of Homer and Russ). This may vary for different users depending on their alveolar size in the histological sections. Click **OK.** **Launch Alveolar Parameters-Multiple Images** To do so, go to: **Plugins ➔ Chord Length Tool Suite ➔ Alveolar Parameters-Multiple Images** (*abbreviated as Alveolar Params-Single Image)* (**Fig. 2g**) Plugin the input parameters (as described above) with the new addition as below: **Minimum alveoli area in calibrated unit**: 25 (the smallest area of each region recognized by the program to be considered a single alveolus. Anything smaller than this number will be filtered out and will not be used for analysis). We did not keep it as ‘0’ because we did not want any artifacts of the alveoli to be counted. **Maximum alveoli area in calibrated unit**: After visualizing several alveoli, we entered this with a value of 100000 so that the algorithm does not go beyond this number which will be unusual for any given alveolus, be it for mouse or humans. **Radius of minimum filter for splitting**: 2 (tuning it will either increase or decrease the radius of the alveoli). Increasing the number will shrink each alveolus and increase the space between adjacent alveoli. The number should be such that the alveolar walls gently touch each other without any overlapping, which might happen if the radius is decreased. **Keep alveoli on the edge:** keep this box □ unchecked. Checking this box will analyze those alveoli which are on the edges of the image frame and unchecking it will eliminate those images which are partially within the frame of the image. Click **OK**. All input information is now saved.

**Fig. 2:**
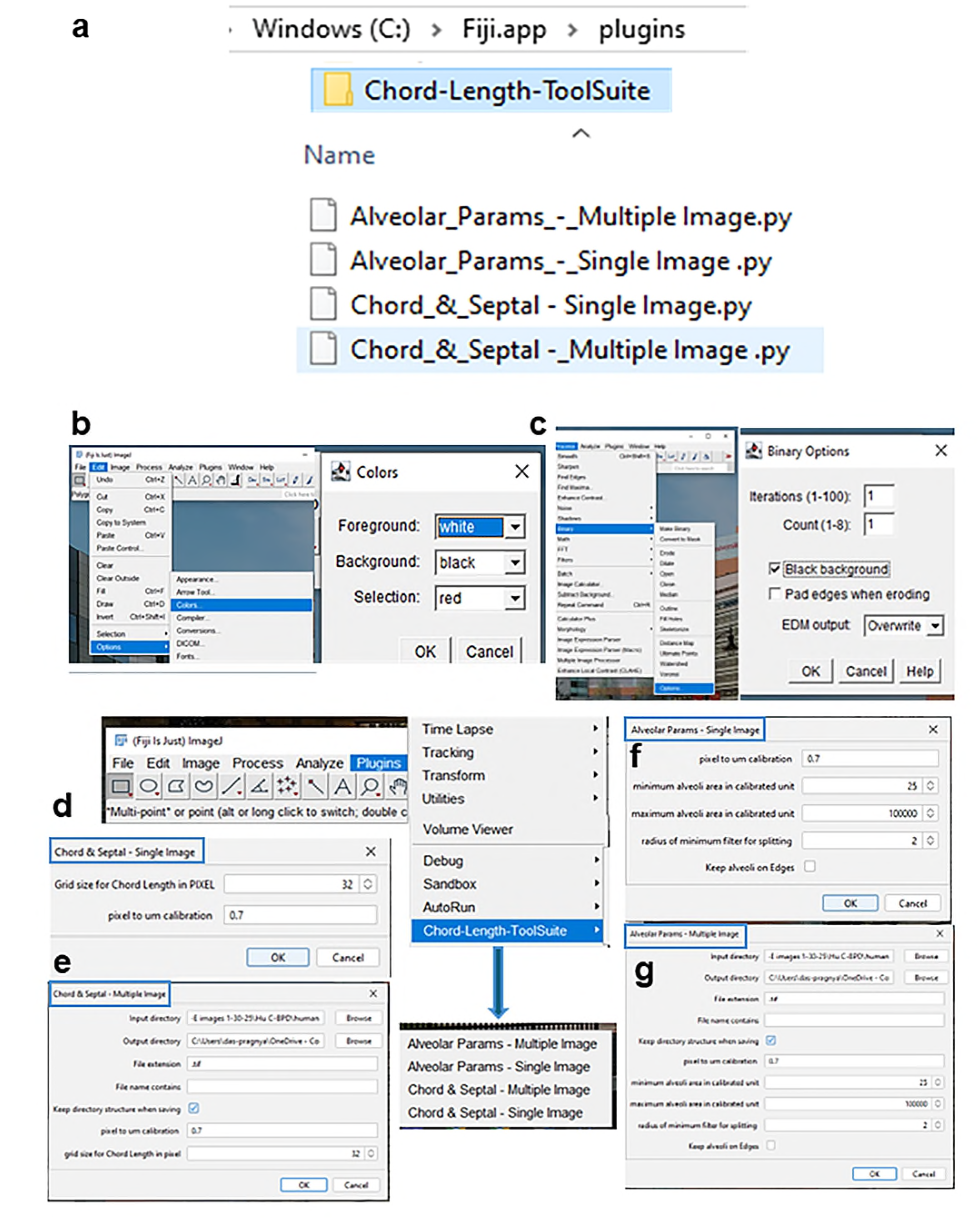
The Chord Length Toolsuite is **(a)** installed under Chord Length plugin of Fiji. With the required image open in Fiji, a single click on any of these macros will analyze either the L_m_ and S_T_ values or other alveolar parameters on the same image. If batch images of the same parameters are to be analyzed, browsing and selecting the required folder will generate the required data. Prior to installing the plugin, the color settings for Chord Length should be made to **(b)** white foreground with black background with the **(c)** binary option of black background. The program has been written in such a way that it automatically inverses the color and binary option for septal thickness while adjusting the color and binary options for chord length; **(d)** shows the options to analyze either L_m_-S_T_ single image in blue rectangle or **(e)** multiple images in blue rectangle **(f -g)** shows the option to analyze other alveolar parameters either for **(f)** single image in blue rectangle or **(g)** multiple images in blue rectangle. The blue solid arrows show the different chord length plugins under the Chord Length ToolSuite. **L_m_:** Chord length; **S_T_:** Septal thickness.

### How the Chord Length & Septal Thickness plugin works: simplified workflow

To measure the alveolar L_m_ and S_T_, we modified and implemented two fully automated methods (Homer, unpublished): (i) one based on grid approach to calculate **Grid Chord Length** and (ii) the second approach based on distance transform known as **Local Thickness Chord Length.** Both these plugins are included under Chord Length macros Folder and are automatically downloaded while downloading the Chord Length ToolSuite. When the macros are run through the lung images, the images are segmented for Grid Analysis (as shown in **Figs. S4A-(a-c)** and Thickness Analysis (as shown in **Figs. S4A (a, d-e)** to extract the said parameters. The results are then automatically combined and presented in spreadsheets in the final form (**Fig. S4A-f**)

#### (i) Grid Chord Length (GCL)

The GCL method is adapted [31] and implemented as an ImageJ Macro [4]. Because L_m_ is defined as the average length of line segments (chords) intersecting the lung mask with a grid, two sets of parallel lines (horizontal and vertical) spaced within a specific grid width (*32 for the present study, as described above*) are overlaid to the binary mask. The binary mask feature is achieved on the lung images using the ImageJ Auto-Threshold Method (https://imagej.net/ij/docs/faqs.html#auto). The intersection points are extracted and their size is then averaged to measure the average GCL.

#### (ii) Septal Thickness

The Septal Thickness is then estimated as the mean of the Local Thickness map of the “inverse” of the binary mask obtained to calculate the Grid Chord Length. The Local Thickness map is extracted with the Local Thickness Fiji Plugin, pre-embedded within the program. The work flow to measure S_T_ is similar to that of L_m_ (**Fig. S4B**), except that the alveoli binary mask used for GCL estimation is ‘inverted’ to identify the septal region. The S_T_ is then estimated as the mean of the Local Thickness map extracted with a similar plugin embedded within Fiji (https://imagej.net/plugins/local-thickness) (**Fig. S4B a-d)**.

#### (iii) Other Alveolar Parameters

The method to calculate the single morphological alveoli parameters is implemented as a Fiji Jython plugin [4]. After converting the images to 8-bit gray scale image, the lung sections are extracted using the Fiji Triangle Thresholding method schema (https://imagej.net/plugins/auto-threshold#triangle) followed by calculating L_m_ thickness map for each lung region using the Fiji Local Thickness algorithm (https://imagej.net/plugins/local-thickness). For each alveolar chord region, different L_m_ estimates (mean, median, mode) are measured from the Local Thickness map, while from the thresholder image, the maximum feret length (distance between 2 parallel planes) and the major and minor axes of a fitted ellipse are measured. Other features like alveolar circularity, area and aspect ratio are also measured automatically using this feature. The macro processes the image first, using an interactive threshold setting, then opens the test grid and subjects the image to a mathematical form to generate lines equivalent to chord length. The program then averages the length. The other macros calculate the length of the interface of alveoli to air. They employ a procedure (called "onion") which allows the Image to see interior holes. Finally, both LTCL and GCL methods are incorporated into a batch analysis in Fiji Jython script to automatically analyze multiple images and collect averaged measurements per field, as described below. The details of grid analysis workflow have already been described earlier.

## Results

The results of the analysis of all images from different groups are included in the output folder. In the subfolders there is the Local Thickness Map of each image. The L_m_ spreadsheet (in two formats .csv and .xls) contains the result of the analysis highlighting showing area, perimeter, circularity, solidity, Maximum Feret Length, Minor and Major axis of fitted Ellipse and Aspect Ratio as defined in the ImageJ documentation [40]. We have automated the following macros in the present work:

a. <u>Chord Length & Septal Thickness - Single Image</u> (will analyze *both* L_m_ and S_T_ of any given single individual image)
b. <u>Chord Length & Septal Thickness - Multiple Image</u> (will analyze L_m_ and S_T_ of multiple images in a single run)
c. <u>Other Alveolar Parameters - Single Image</u> (will analyze other parameters like alveolar area, perimeter, circularity, solidity, major axis, minor axis and thickness of a given single individual image)
d. <u>Other Alveolar Parameters - Multiple Image</u> (will analyze other parameters like alveolar area, perimeter, circularity, solidity, major axis, minor axis and thickness of images in large batches of multiple files, simultaneously)

### Adding shortcut key to the plugin

To hasten the image analysis, Fiji allows the use of shortcut keys by bypassing multiple clicks on the specific plugin every time. Users should refer to the use of Fiji manual to add this feature [40, 41]. Here, we have added two shortcut keys – ‘6’ (to measure L_m_ and S_T_-Single Image) and ‘7’ (to measure Other Alveolar Parameters-Single Image), automatically (**Fig. S5** and **SF3**).

### Validation of the plugin

We now have a fully functional Chord Length Plugin with an additional feature of Septal Thickness plugin (https://doi.org/10.5281/zenodo.15354633). The users can access this DOI, run their images through this dual plugin feature and get the desired values **(SF4)**. Let us validate this plugin by a simple case study with 3 images –**(a)** from mouse RA group P14, **(b)** from mouse BPD group P14 (where the lungs are subjected to injury till P4 and then allowed for a recovery phase till P14), **(c)** from hyperoxia-induced acute lung injury (HALI) group P7, where mouse lungs are subjected to injury for 7 consecutive days without any recovery.

1. Launch Fiji
2. Open your image folder; drag a single image (Case **a,b,c**) from your input folder and drop it on the Fiji tab rectangular box ▭
3. Go to **plugins➝ Chord Length ToolSuite ➝ Chord Length & Septal Thickness-Single Image ➝ OK** (To use the shortcut key here: after step 2**, press ‘6’ on the keyboard** ➝ **OK)**
4. The L_m_ and S_T_ values are displayed for the above 3 groups (**Fig. 3a-c**)

**Fig. 3:**
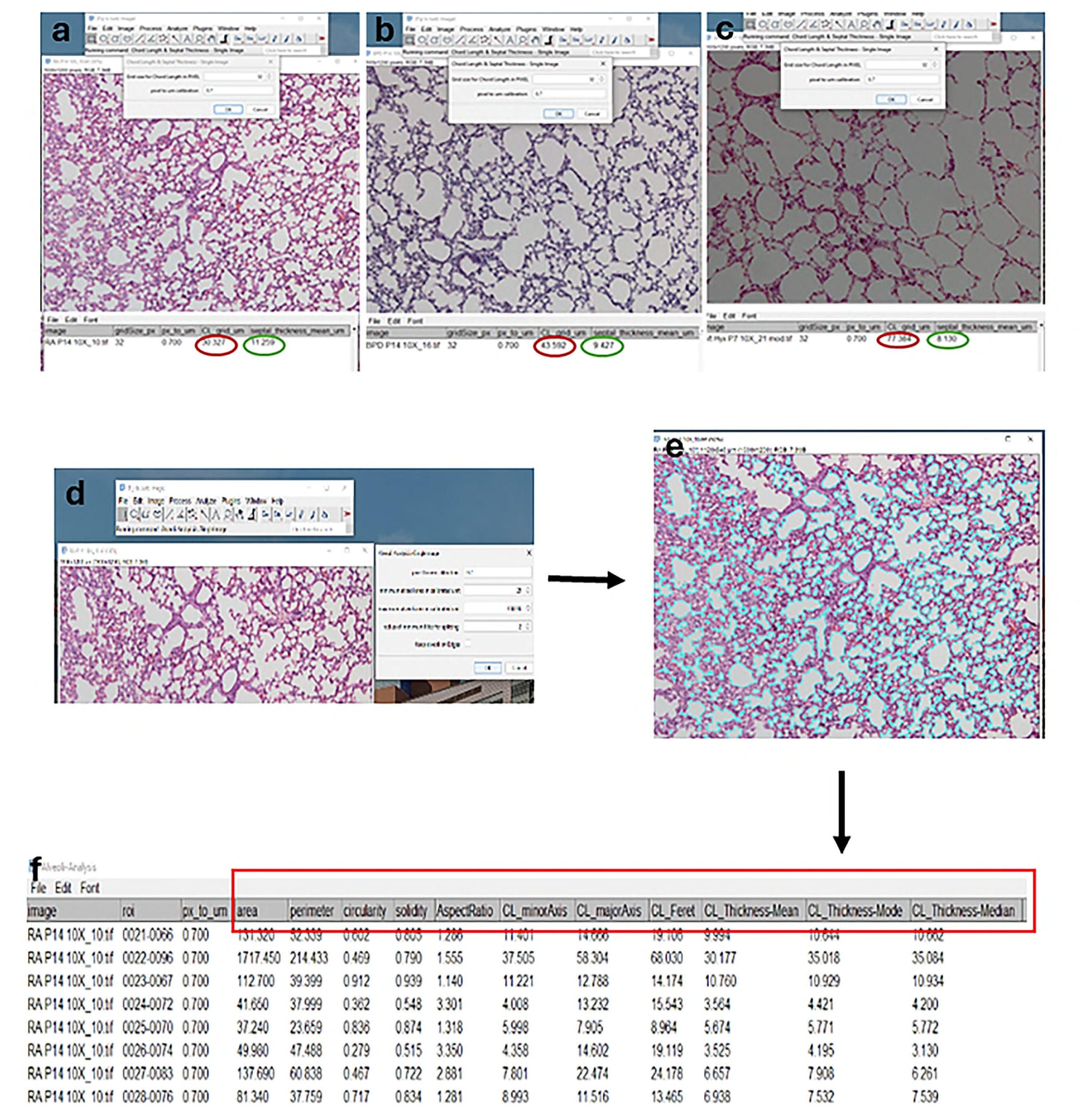
Validation of the plugin to measure L_m_, S_T_ and other alveolar parameters in **(a)** P14 RA neonatal mouse pup **(b)** P14 BPD neonatal mouse pup **(c)** P7 HALI neonatal mouse pup. The plugin measures L_m_ as **30.3**µm**, 43.6**µm**, 77.6** µm and S_T_ as **11.25**µm, **9.42**µm and **8.13**µm for RA, BPD and HALI group, respectively, thus confirming the morphological changes in the alveoli in 3 different conditions. The red circles in “**a,b,c”** denote L_m_ values while the green circles depict S_T_ values; the red rectangular box in “**f”** denotes the different alveolar parameters. Using the dual Chord Length-Septal Thickness Single Image plugin, the algorithm can measure both the parameters simultaneously (red and green circles). **P:** postnatal day; **L_m_:** Chord length; **S_T_:** Septal thickness; **RA:** Room air; **BPD:** Bronchopulmonary dysplasia; **HALI:** Hyperoxia-induced lung injury.

To measure other parameters like alveolar thickness, area, perimeter etc. (with the image already open):

1. Go to **plugins ➝ Chord Length ToolSuite ➝ Other Alveolar Parameters - Single Image ➝ OK** (To use the shortcut key for the above parameter**, press ‘7’ on the keyboard (***with the image still open) ➝* **OK**
2. Multiple parameters are displayed (**Fig. 3f**). Basically, the plugin analyzes every single closed alveolus (as seen in **Fig. 3e**) on the open image and calculates the required parameters, when run through the macros. In sum, with a single image open, L_m_, S_T_ and other alveolar parameters can be measured simultaneously.

Additionally, we also tested lungs of neonatal higher mammals like rabbit [42], lamb [43], baboons [44] (including some unpublished data) and humans [45] born prematurely under different conditions to assess their respective L_m_ and S_T_, using our dual morphometry plugin (**Figs. 4a-e; i-iii**). The plugin was able to quantify the difference in alveolar morphology in all experimental conditions. We can now assert that our customized Chord Length and Septal Thickness Fiji plugin can successfully measure the L_m_ and S_T_ of any given newborn lung with just a click of the mouse, without having to do the additional work of drawing lines manually, counting the intercepts that cross the alveoli and then using a formula to calculate the average MLI.

**Fig. 4:**
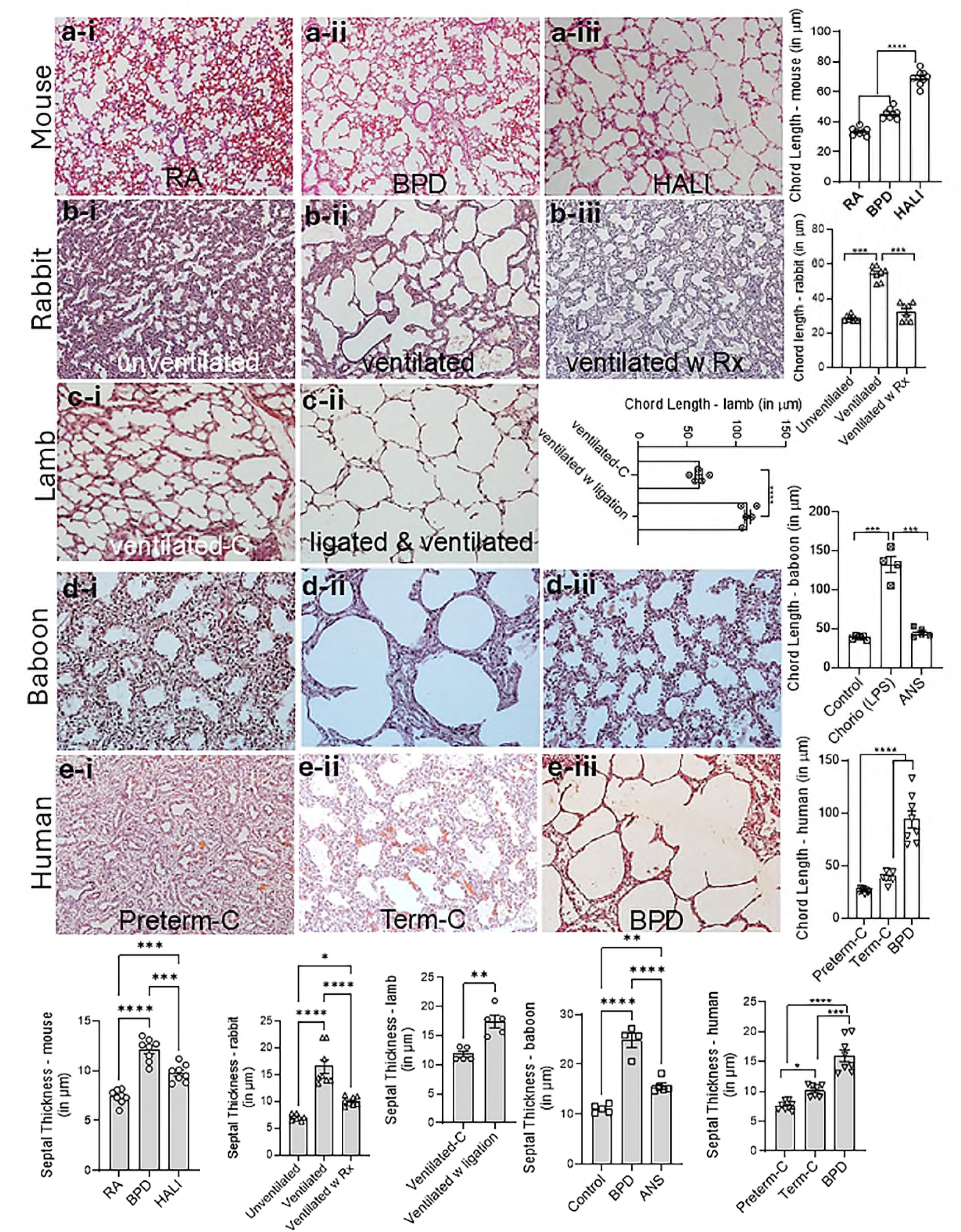
Validating and measuring L_m_ and S_T_ using our custom-made plugin in **(a)** mice **(b)** rabbit **(c)** lamb **(d)** baboon and **(e)** human neonates. In mice, (**a-ii**) newborn pups were either exposed to hyperoxia from P0-P4 and allowed to recover in RA till P14 to mimic human BPD or (**a-iii**) exposed to hyperoxia for 7 consecutive days from P0-P7 to mimic HALI. Note the significant enlargement of the alveoli (L_m_) and increased S_T_ in HALI and BPD as compared to RA (room air) controls. Interestingly, there was significantly decreased S_T_ in the HALI model, compared to the BPD one. (N=6-8, males and females combined). Similarly in newborn rabbit kits, the (**b-ii**) L_m_ and S_T_ increased significantly after ventilation when compared to (**b-i**) controls born preterm. When the kits were (**b-iii**) given exogenous surfactant (Curosurf®) and ventilated, there was a significant improvement in the L_m_ and S_T_ (N= 8). In newborn lambs delivered preterm by c-section, followed by ligating the ductus arteriosus to induce pulmonary hypertension, the L_m_ and S_T_ was increased significantly (**c-ii**) in this group compared to the (**c-i**) non-ligated but ventilated group (N=5). In the baboon preterm model (**d-i**; 125d gestational control) when the mothers were administered antenatal steroids and lipopolysaccharide (LPS) during pregnancy to mimic chorioamnionitis, the (**d-ii**) babies (antenatal LPS only; 125d gestation) had a lung phenotype similar to BPD, characterized by marked enlargement of airspaces indicated by increased L_m_ and S_T_ as compared to gestational controls. In (**d-iii;** 125d gestation) the mother was treated with antenatal steroids only, the L_m_ was improved compared to the chorioamnionitis model, but was similar to the gestational control; however, the S_T_ was significantly improved compared to both the gestational control and the chorioamnionitis group (N=4-5). In human babies diagnosed with (**e-iii**) established BPD, the autopsy lung sections show marked enlargement of the alveoli with a significant increase in L_m_ and S_T_ compared to (**e-ii**) lungs of babies born full term with no respiratory complications and (**e-i**) lungs of babies born preterm. (N=7-8; males and females combined). The bottom panel shows S_T_ in all the models. **\*\*\***P<0.005; **\*\*\***P<0.0001. **P:** postnatal day; **L_m_:** Chord length; **S_T_:** Septal thickness; **RA:** Room air; **BPD:** Bronchopulmonary dysplasia; **HALI:** Hyperoxia-induced lung injury; **Rx:** treatment with exogenous surfactant (Curosurf®); **C:** Control; **w:** With; **Chorio:** Chorioamnionitis; **LPS:** Lipopolysaccharide; **ANS:** Antenatal Steroids

### Problems, Limitations and Conclusions

In this review we have modified a simple custom-made pre-existing Fiji plugin which can automatically measure the alveolar L_m_ and S_T_ of both small and large experimental animals whose lungs were injured or damaged due to some form of insult (including autopsied neonatal human lungs who died of BPD) as well as those whose lungs recovered after treatment in experimental BPD models. Sometimes when analyzing large batches of images, the program might crash due to excessive data overload. In such a case, users should consider exiting the program and rebooting the PC again. If the program is downloaded in the Institution’s core computer, the program might be deleted at times during regular clean-up of the PCs, and users must reload it by downloading it again from the link that we have provided, or from their backup site. Every time a plugin or a macro is modified or edited, the application needs to be restarted. The only limitation in this study is that we have edited a preexisting publicly available macro and have not described the principles on which the macro is based which is beyond the scope of the present work, but at the same time we wanted uniformity in interpretation because different laboratories employ different methods for measuring various parameters of the lung. Our aim is to introduce this concept and share the plugin with all investigators employed in pulmonary research to get uniform data without having to invest heavily in sophisticated stereology software. Our modification is not based on any mathematical validation of the algorithm. We have made the plugin user-friendly for easy accessibility and practicality so that any individual can use this plugin for their research purpose. After validating our plugin in different animal models, users can employ it to measure similar functions taking into account different considerations that are discussed in the beginning. At the same time, we do not advise users to rely solely on our L_m_ - S_T_ measurement method as the “only” method to draw conclusions about a study and should depend on additional supporting experiments/measurements to back up their findings. Although there is a lot of literature that explains automating the plugins and introducing new features to measure L_m_, our goal was not to overwhelm readers with the technology but to introduce a simple solution to measure an important readout as a parallel surrogate with the aligned research study.

In the end, we also want to make the young readers aware of the charitable contribution of Ewald R Weibel, Domingo M. Gomez, Wayne Rasband and John C Russ who laid the foundation for morphometry in Pulmonology and automated the quantification by developing the ImageJ software for free public use. This work of ours is a humble tribute to the magnanimous contributions of these stalwarts in morphometry and image processing.

### Points for clinical practice and questions for future research

1. We have validated our custom-made plugin in human lung sections so that it can be used in the clinical setting to measure L_m_ on autopsy or biopsy sections, if needed.
2. So far, L_m_ has not been measured in higher animals due to the non-availability of a specific automated plugin. We have made our plugin freely available for all scientific users.
3. The original scripts for the macros are also available as supplementary material which can be modified for personalized use depending on the needs of a particular researcher.
4. Future work should consider using other new morphometric standards along with L_m_ and S_T_.

## Acknowledgements

We would like to thank Fabrizio Salomone, Costanza Casiraghi, and Nicola Pelizzi (Chiesi Farmaceutici, Italy) for the preterm rabbit slides, Satyan Lakshminrusimha and Sylvia Gugino (University of Buffalo, NY, USA) for the preterm lamb slides, Shamim Mustafa and Steve Seidner (UT Health, San Antonio, Texas, USA) for the preterm baboon slides and Sture Andersson (Helsinki, Finland) for the human slides, respectively.

## Author Contributions

**Conceived by:** PD

**Data Acquisition:** PD, EM, OA, ET, DTR, BA, RJH

**Funding:** PD, EM

**Manuscript draft:** PD, EM

**Manuscript Editing:** PD, EM, BA, RJH, VB

All authors have approved the final version of this manuscript.

## Supplemental Figure Legends

**Fig. S1:**
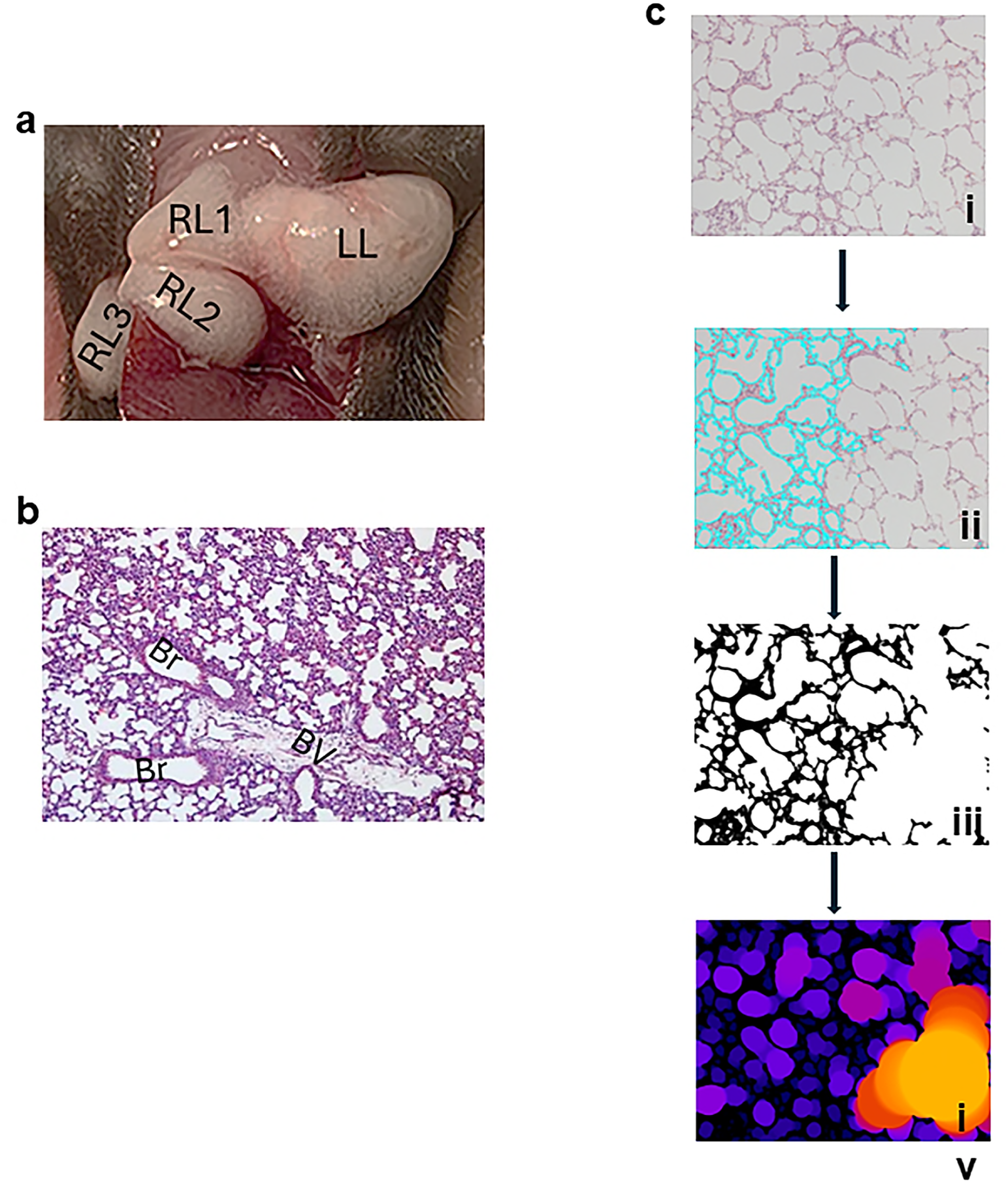
(a) Harvesting of clean lung tissue from P14 mouse pups for histological sectioning.; **LL:** left lung; **RL1-3:** right lung lobes. RL4 is hidden below RL1-3; **(b)** Histological section of P14 mouse lungs showing bronchioles (Br) and large blood vessels (BV) that may interfere in measuring the chord length; **(c)** faint hematoxylin-eosin staining **(i)** on the lung sections to show that (**ii**) overlay and **(iii)** binary conversion will be affected before calculating the **(iv)** final L_m_ and S_T_ readout. The empty space in **(iii)** will result in a blob in **(iv)**. **P:** postnatal day; **L_m_:** Chord length; **S_T_:** Septal thickness.

**Fig. S2:**
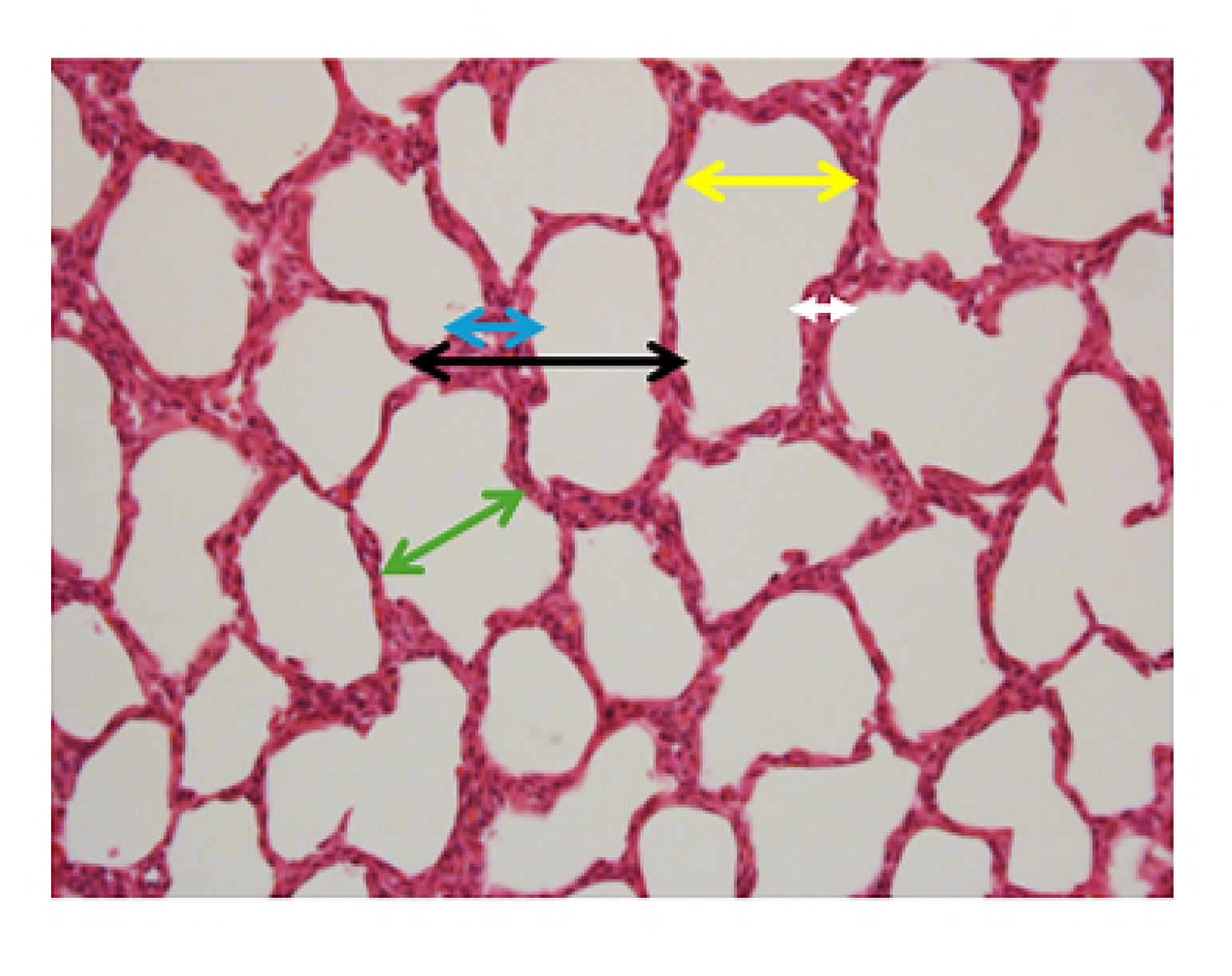
Histological section of P14 mouse lungs showing different alveolar parameters measured for morphometry. Black double-headed arrow denotes mean linear intercept (MLI); yellow arrow shows chord length, green arrow depicts alveolar diameter, white arrow-alveolar wall, blue arrow – septal thickness. **P:** postnatal day.

**Fig. S3:**
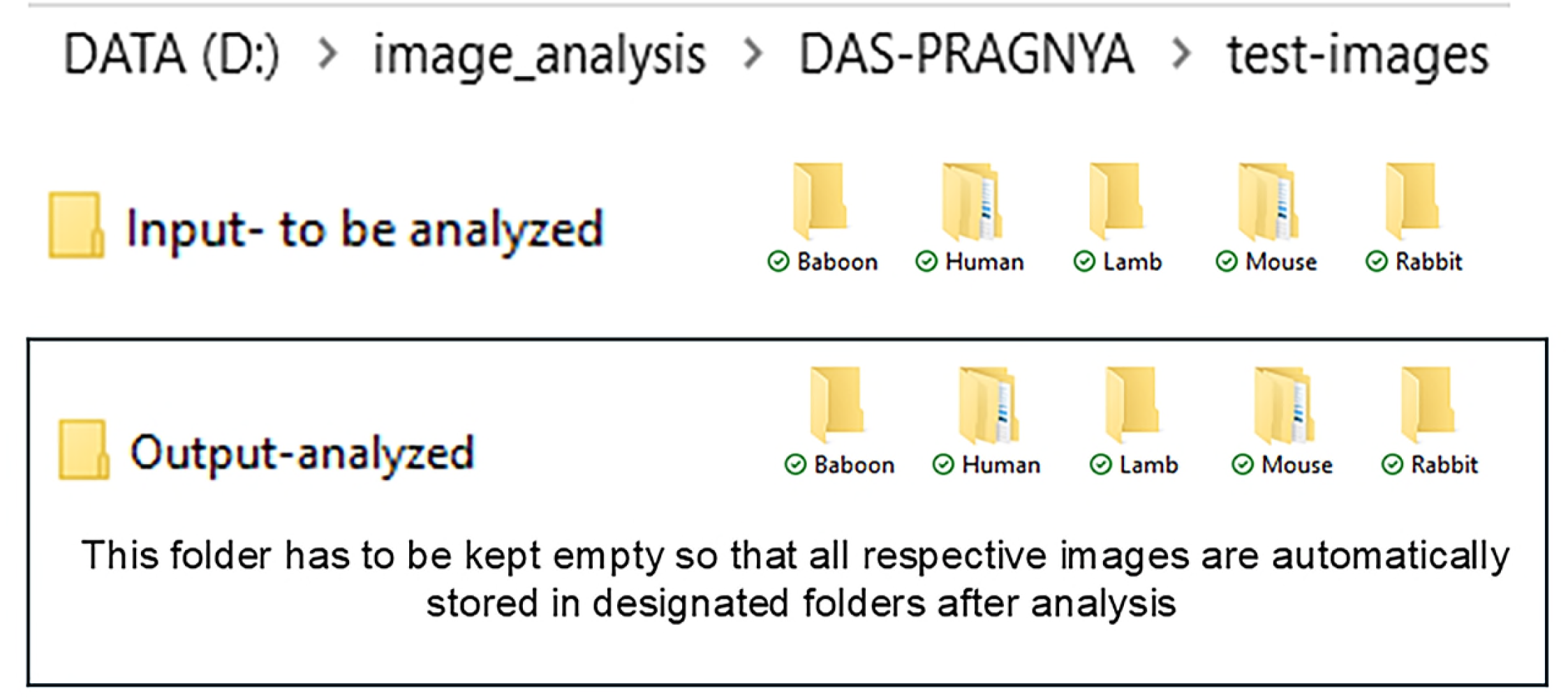
While analyzing multiple images using the batch analysis plugin, all the images should be placed in the appropriate folder (input and output) so that the browsing option (as shown in **Figs. 3b, d**) will help to upload all the images from the input folder simultaneously and analyze the data in a single attempt. The data analyzed will then be automatically stored in the output folder. Hence, prior organization is necessary for this systematic profiling of the analyzed values.

**Fig. S4:**
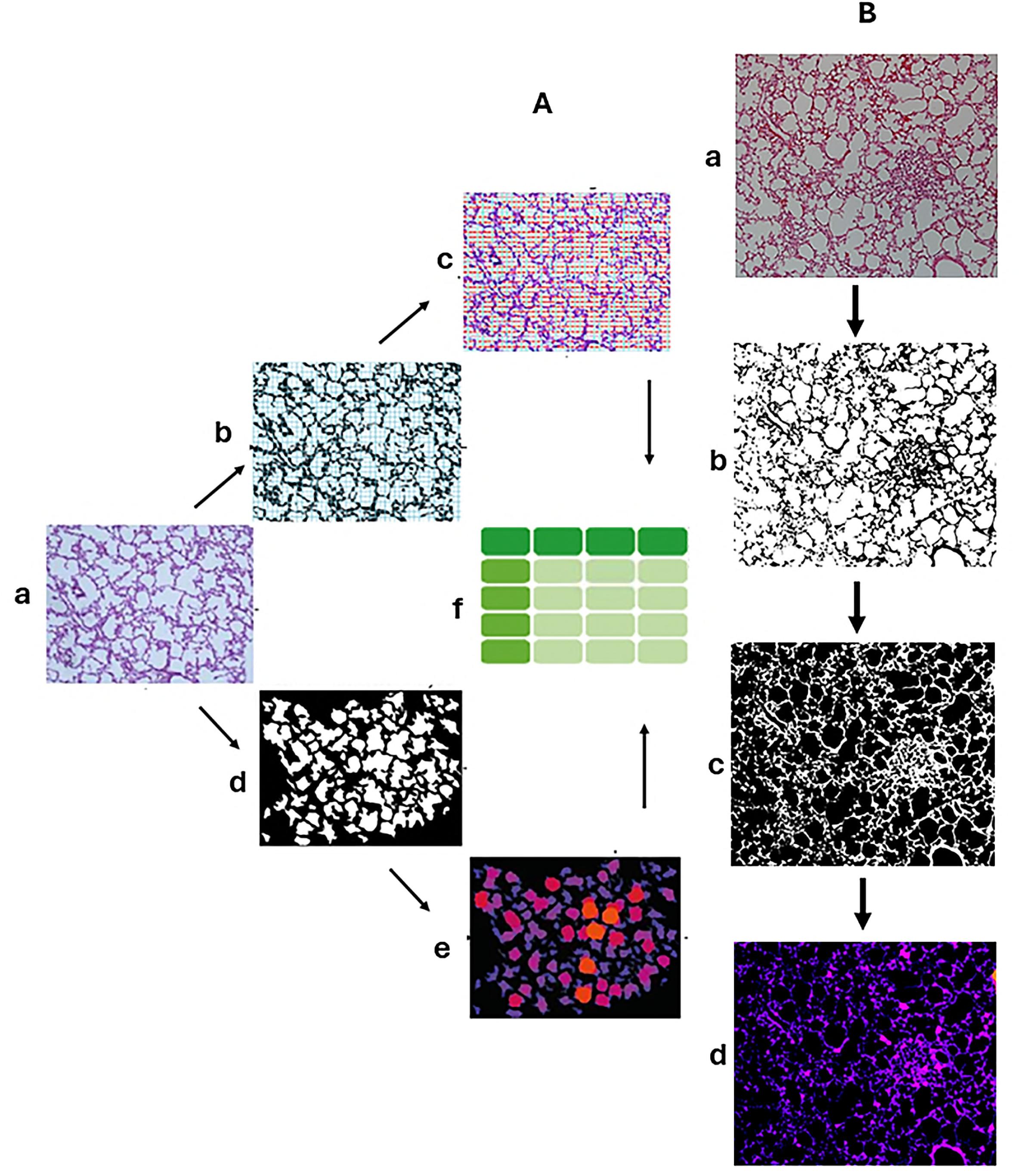
Workflow of the **(A)** Chord Length plugin installation process showing the masking of **(a)** original lung images in **(b)** binary mode and extracting the chord Length values by **(c)** automatically drawing vertical and horizontal grids over the alveolar sacs; **(d)** masking of original lung images in **(d)** binary mode and analyzing the **(e)** thickness of each alveolar sac. The algorithm of the Chord Length Plugin gives a cumulative readout of the total alveolar grid length and total thickness, which is presented in **(f)** as a combined feature of the results in .xls format. Similarly, to use **(B)** Septal Thickness plugin, the original **(a)** H-E images are **(b)** converted to binary images and then **(c)** extracted by inverting the images to result in **(d)** local Septal Thickness map.

**Fig. S5:**
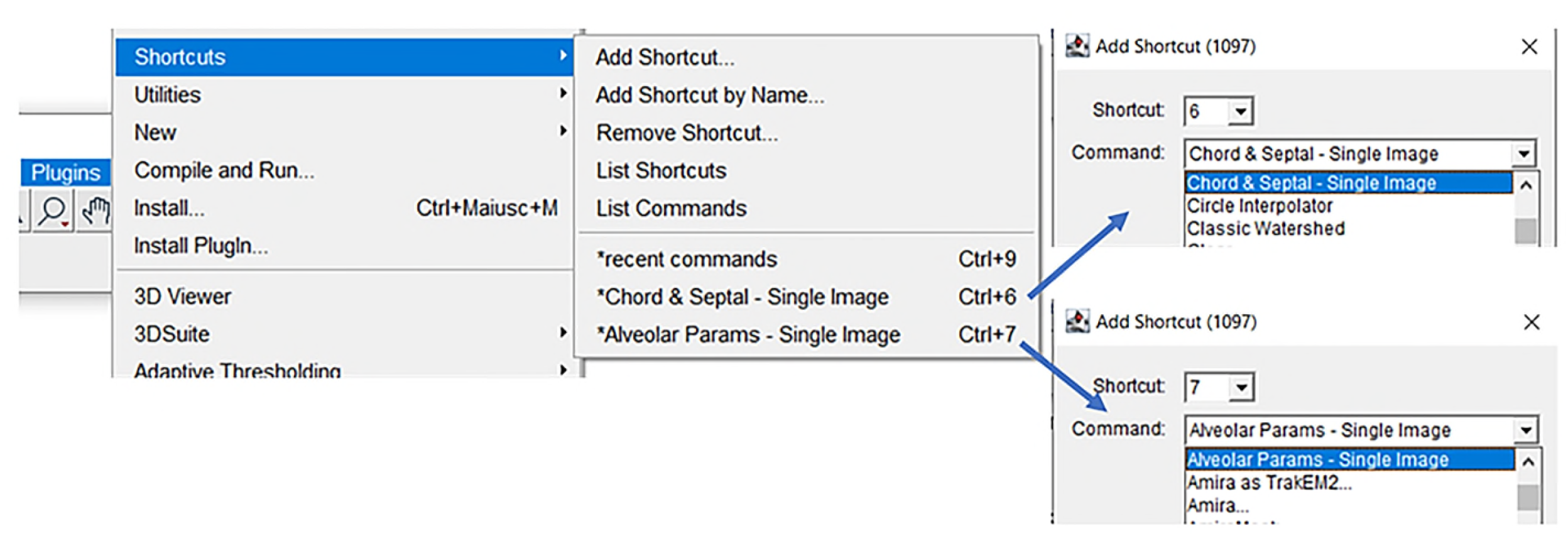
Image analysis can be expedited by **(a)** adding shortcut keys to the plugin. A shortcut key of **(b)** ‘6’ or ‘ctrl+6’ will automatically analyze L_m_ and S_T_ while a shortcut key of **(c**) ‘7’ or ‘ctrl+7’ will automatically analyze other alveolar parameters on lung sections, for a given single image. **L_m_:** Chord length; **S_T_:** Septal thickness.

## Supplemental File 1

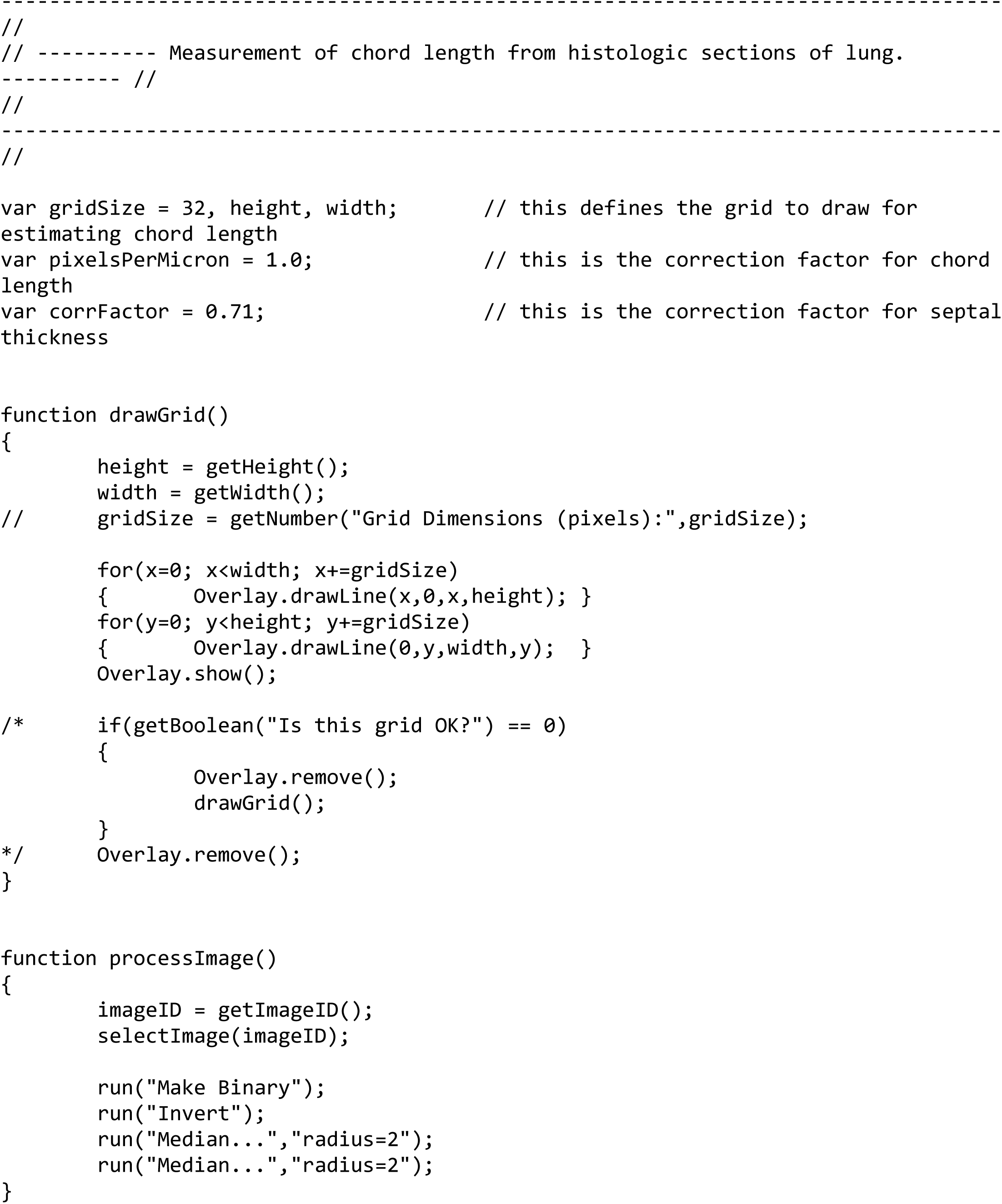
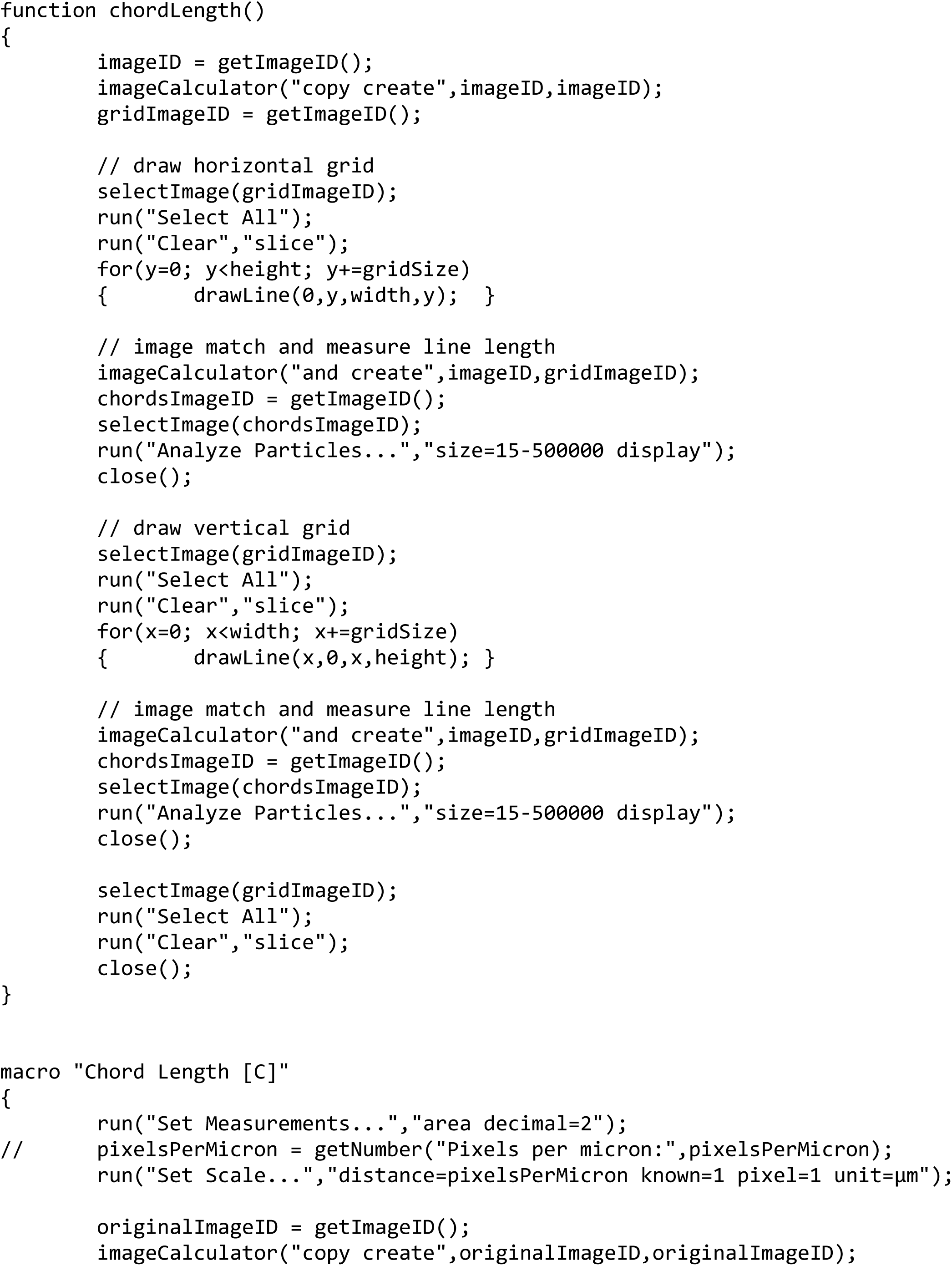
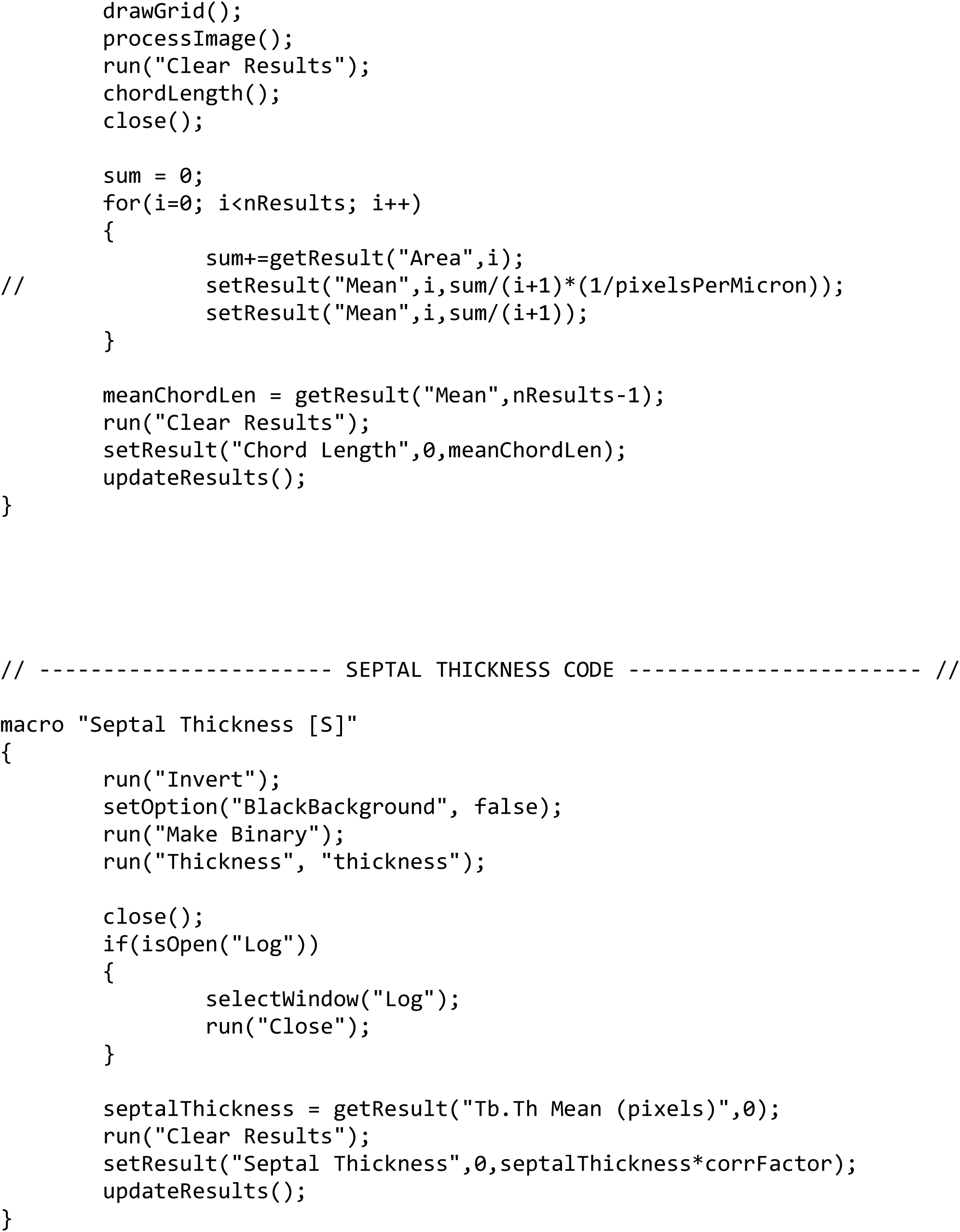

## Supplemental File 2

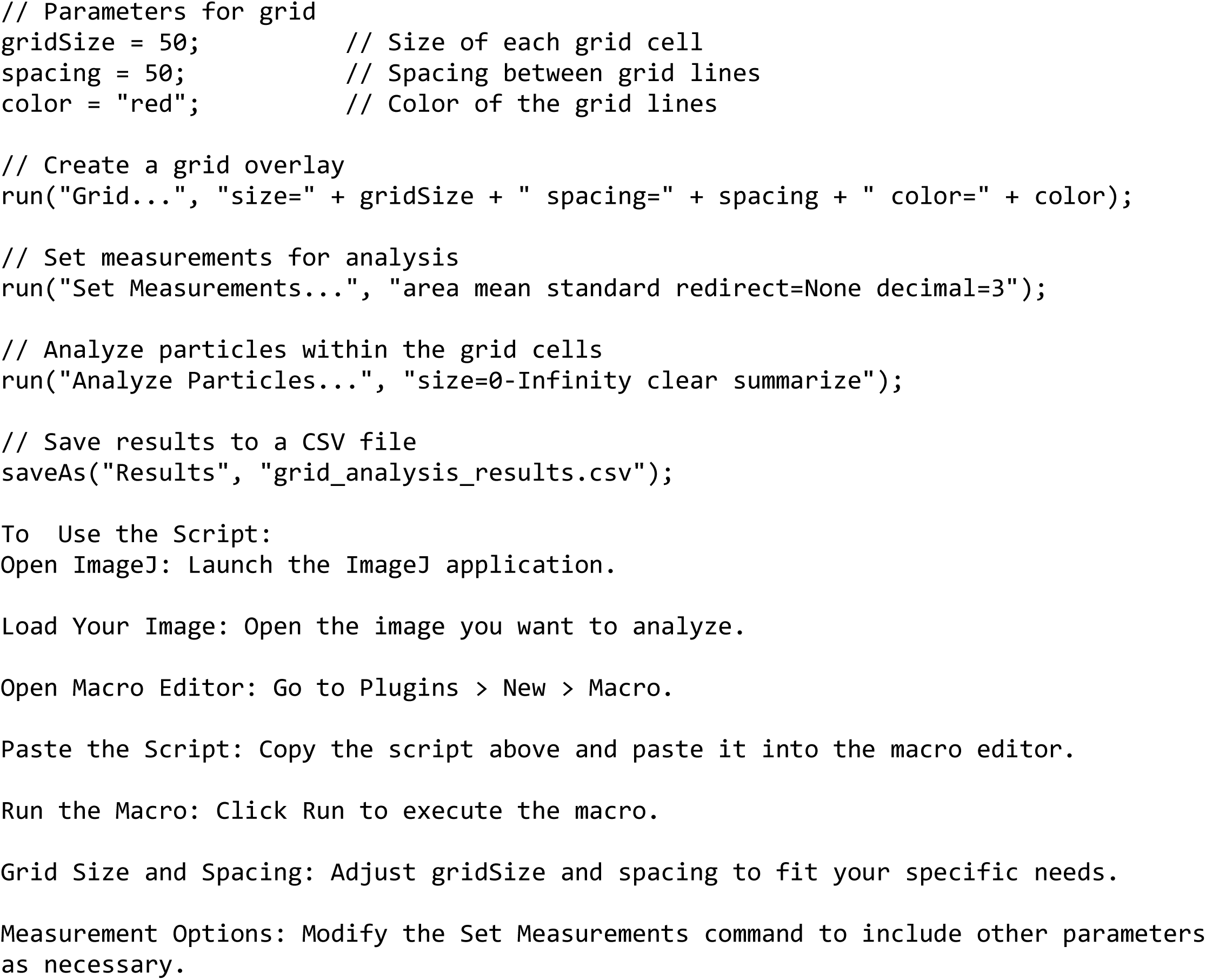

## Supplemental File 3 (SF3)

**Adding shortcut keys for Chord Length & Septal Thickness-Single Image and <u>Other Alveolar parameters - Single Image</u> analysis (Fig. S5 a-c)**

Fiji allows its users to assign a shortcut key to launch a specific plugin (https://imagej.net/ij/docs/guide/146-31.html#toc-Subsection-31.2) so as to fasten the analysis by avoiding to click on the specific plugin every time an image is opened for analysis. To do so, go to:

1. **Plugins** ➔ **Shortcuts** ➔ **Add Shortcut** (a pop-up window appears)
2. From the dropdown **Shortcut** menu, select any number or alphabet etc. to assign a shortcut key. In this study, we have added ‘6’. (*The users should keep in mind that the drop-down list will differ for every user depending on what key has already been used previously, and a common list will not be available to every user*).
3. From the dropdown **Command** menu, search for ‘Chord & Septal Single Image’
4. Click **OK**.

The shortcut key ‘6’ is assigned to **Chord Length & Septal Length - Single Image Analysis**

Repeat steps 1 through 4 to assign another shortcut key to **Other Alveolar Parameters – Single Image**. In this study, we have added ‘7’ as the shortcut key for this plugin.

To validate the shortcut keys:

1. Lauch Fiji
2. Open your image folder; drag and drop a single image from your image folder ▭
3. Press **‘6’** on the keypad

The pop-up window appears. Click **OK**

The **Chord Length & Septal Length measurements** are displayed automatically. Similarly,

4. Press **‘7’** on the keypad with the same image open as above.

The pop-up window appears. Click **OK**

The **Other Alveolar Parameters** are displayed automatically.

The readouts of the above 2 parameters will be generated in .csv format which can be exported as .xls format. For the present work, we were more focused on measuring L_m_ and S_T_ rather than all the additional alveolar parameters.

## Supplemental File 4 (SF4)

**Instructions to download the plugin from the DOI link (**https://doi.org/10.5281/zenodo.15354633)

1. Download the Fiji plugin on your PC as described in the text with all its associated files.
2. Click on the DOI link or copy and paste the link on a browser to download the Chord Length ToolSuite.
3. Extract the zipped ToolSuite file and move the extracted file folder to the location where Fiji plugin is installed.
4. All files are now placed in one location and ready to use.

